# Structure and regulation of the myotonic dystrophy kinase-related Cdc42-binding kinase

**DOI:** 10.1101/2022.03.11.483953

**Authors:** Linda Truebestein, Elisabeth Waltenberger, Charlotte Gehin, Anne-Claude Gavin, Thomas A. Leonard

## Abstract

Remodeling of the cytoskeleton underlies myriad processes essential for life. Protein kinases of the DMPK family are critical regulators of actomyosin contractility in cells. In the nematode worm, *Caenorhabditis elegans*, MRCK1 is required for the activation of myosin, leading to the development of cortical tension, apical constriction and early gastrulation. Here, we present the structure, conformation, and membrane-binding properties of *C. elegans* MRCK1. MRCK1 forms an obligate homodimer with N-terminal kinase domains, a parallel coiled-coil of 55 nm, and a C-terminal tripartite module of C1, PH and CNH domains. High-throughput liposome binding assays indicate binding to specific phosphoinositides, while the C-terminal Cdc42-binding (CRIB) motif binds specifically to activated Cdc42. The length of the coiled-coil domain of MRCK, as well as those of the related DMPK kinases ROCK, CRIK and DMPK, is remarkably conserved over millions of years of evolution, suggesting that they may function as molecular rulers to precisely position kinase activity at a fixed distance from the membrane.

## Introduction

Cell shape remodeling and cell movement are necessary for physiological processes such as fertilization, cell division, and the morphogenesis and motility of intracellular organelles, as well as pathophysiological processes such as the phagocytosis of invading pathogens, cancer cell invasion, and metastasis. These changes in cell shape and motility are accomplished primarily by the contraction of actomyosin filaments within the actin cytoskeleton (1).

In order for cells to adopt defined shapes and be capable of directed movement, the contractile machinery must be exquisitely regulated. At the heart of this regulation is the control of myosin II activity by phosphorylation of its regulatory light chain (RMLC/MLC2). Phosphorylation of two conserved residues in the N-terminus of MLC2 stimulates the ATPase activity of myosin, leading to actomyosin contraction (2). MLC2 is phosphorylated by myosin light chain kinase (MLCK) (3) and dephosphorylated by myosin light chain phosphatase (MLCP) (4–6).

The dystonia myotonica protein kinase (DMPK) family, comprising Citron Rho-interacting kinase (CRIK), myotonic dystrophy protein kinase (DMPK), myotonic dystrophy kinase-related Cdc42-binding kinase (MRCK), and Rho-associated coiled-coil kinase (ROCK), are known regulators of MLC2 phosphorylation. *In vitro,* ROCK and MRCK are capable of directly phosphorylating MLC2 (7–9), but primarily exert their effects *in vivo* via the inhibitory phosphorylation of MLCP (10–12). As such, they play key roles in cytoskeletal remodeling and have been implicated in diverse cellular processes from cell protrusion (9), neurite outgrowth (13) and cytokinesis (14) to embryonic elongation (15, 16), nuclear positioning (17) and establishment of epithelial polarity (18, 19). Triplet expansion of the DMPK1 gene is causative of autosomal dominant muscular dystrophy 1 (20–22), while ROCK and MRCK have been shown to play essential roles in neural tube closure during embryonic development (23), as well as cancer cell invasion and metastasis (10, 24, 25).

The DMPK kinases are members of the AGC kinase family with a conserved domain architecture of N-terminal kinase domains followed by a predicted coiled-coil domain and C-terminal regulatory domains that mediate a variety of protein-protein and protein-membrane interactions. While many AGC kinases are regulated by phosphorylation of motifs in their activation loop or C-terminal tail (26), the DMPK kinases are not. Instead, their N-terminal capped-helix bundle (CHB) domain cooperates with the C-terminal tail to maintain the kinase domain in a homodimeric, active conformation with an ordered activation loop (27–29). Plasma membrane localization of ROCK and CRIK is reported to be driven by RhoA (14, 30), while Cdc42 mediates the recruitment of MRCK (9). DMPK1, which is almost exclusively expressed in skeletal and cardiac muscle (31), is presumably anchored to the membrane via its predicted C-terminal transmembrane domain. While a recent study of ROCK2 suggested that its activity is regulated by a combination of its subcellular localization and the length of its coiled-coil domain (7), the mechanisms by which other DMPK kinases are regulated are still largely unknown.

The nematode worm *C. elegans* encodes a single *mrck* gene (mrck-1), with closest homology to human MRCKα. MRCK1 has been shown to be essential for apical constriction during gastrulation in *C. elegans* (15) and is localized to the apical membrane by activated, GTP-bound Cdc42, where it drives actomyosin contractility via increased phosphorylation of MLC2. Conversely, Cdc42 is locally inactivated on the basolateral membrane, thereby inhibiting the recruitment of MRCK. To better understand how MRCK1 is localized and how its activity is regulated, we have determined the low-resolution structure and conformation of full-length MRCK1 as well as the high-resolution structure of its regulatory domains. Using diverse membrane binding assays, we have determined which lipids can be recognized by MRCK1 at the membrane surface. Finally, we have quantified MRCK1 binding to active Cdc42 *in vitro. We* propose a model in which MRCK1 is recruited to the apical membrane by the coincident recognition of membrane lipids and active Cdc42, which serve to position its kinase domains at the end of its coiled-coil domain at a fixed distance from the membrane. Bioinformatic analysis reveals remarkable conservation of the lengths of the coiled-coils of all DMPK kinase family members, thereby extending the concept of the coiled-coil domain as a ‘molecular ruler’ capable of governing protein kinase signaling and actomyosin contraction at precise molecular subcellular locations.

## Results

### Full-length MRCK1 adopts a ROCK-like extended, active conformation

*C. elegans* MRCK1 consists of an N-terminal CHB domain which mediates kinase domain dimerization (28), followed by a region of predicted coiled-coil and C-terminal C1, pleckstrin homology (PH), and citron homology (CNH) domains, of unknown structure and function (Figure 1A). A Cdc42 and Rac-interactive binding (CRIB) motif is encoded C-terminally to the CNH domain. Bioinformatic analysis of the coiled-coil domain revealed that, despite variation in the length of the sequence between the N-terminal kinase domain and C-terminal regulatory domains (Figure 1B), the length of the predicted coiled-coil domain is remarkably well conserved (Supplementary Figure S1A), reminiscent of the strict length conservation in the related ROCK kinases (7, 32). To investigate the structure and conformation of a full-length MRCK protein, we purified recombinant *C. elegans* MRCK1 from baculovirus-infected Sf9 insect cells to homogeneity. MRCK1 exhibited robust kinase activity against recombinant myosin light chain 2 (MLC2), the regulatory subunit of myosin (Figure 1C). Fitting of the data with the Hill equation indicated a Hill coefficient of close to 2, suggesting cooperativity in the phosphorylation of MLC2. Purified MRCK1 was imaged by rotary shadowing electron microscopy, revealing an extended particle, 70-85 nm in length, comprising globular regions of electron density at either end of a long, semi-rigid coiled-coil domain with an average length of 55.0 ± 7.2 nm (Figure 1D). Dimeric ROCK2 particles exhibit comparable barbell-shaped structures with a length of 106.7 ± 3.5 nm (Figure 1E), corresponding to the length of predicted coiled-coil between its N-terminal kinase and C-terminal regulatory domains (7). The length of the coiled-coil domain of ROCK2 is highly conserved over 650 million years of evolution and has been shown to be important for stress fiber formation in cells (7). While the number of intervening amino acids in MRCK1 would predict a canonical, uninterrupted coiled-coil of 78.1 nm, we observed a shorter mean distribution of particle lengths and a bigger fluctuation from the mean than in ROCK2 (Figure 1F). *In silico* prediction of the coiled-coil propensity of the intervening amino acids revealed four segments of coiled-coil with a total predicted length of 54.6 nm (Supplementary Figure S1A). The structural explanation for why fulllength MRCK1 dimers are considerably shorter than expected (taking into account regions of predicted disorder) is not obvious and will likely require further investigation, though the predicted coiled-coil length of ~55 nm (Supplementary Figure S1A) is conserved over 650 million years and corresponds to the mean length observed.

**Figure 1.**
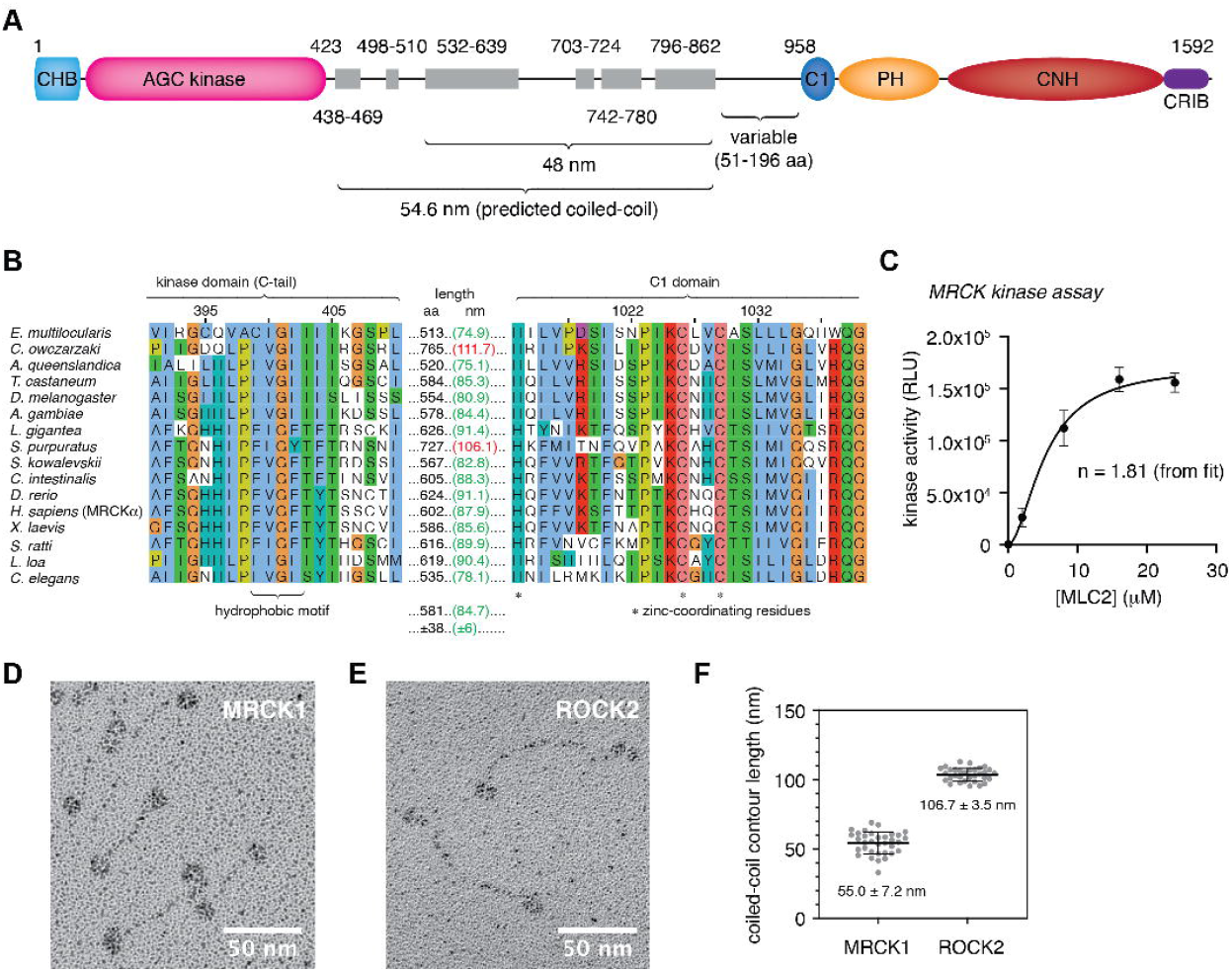
Full-length MRCK1 adopts a ROCK-like extended, active conformation. A. Domain architecture of *C. elegans* MRCK1. (CHB, capped helix bundle domain; C1, protein kinase C conserved domain 1; PH, pleckstrin homology domain; CNH, citron homology domain; CRIB, Cdc42 and Rac-interactive binding domain). B. Partial sequence alignment of MRCK orthologs. The length of the intervening sequence between the end of the kinase domain and the beginning of the C1 domain is indicated as number of amino acids (black) and predicted length of a canonical parallel coiled-coil (green) (assuming continuous parallel coiled-coil with a pitch of 1.48 Å/residue). C. MRCK1 kinase activity against recombinant MLC2. Data fitted with the Hill equation. D. Rotary shadowing electron microscopy of full-length *C. elegans* MRCK1. E. Rotary shadowing electron microscopy of full-length *H. sapiens* ROCK2. F. Contour length distribution of the coiled-coil domain of MRCK1 and ROCK2.

### Structure and properties of the C1-PH-CNH module of MRCK1

The C-terminus of MRCK contains predicted C1, PH and CNH domains of unknown structure and function. To shed light on the function of this regulatory module, we determined the structure of a construct of MRCK1 comprising residues 955-1534 by X-ray crystallography. The protein crystallized in space group P1 with 5 molecules per asymmetric unit and crystals diffracted to 2.14 Å. Efforts to determine the structure either by molecular replacement, multiple isomorphous replacement, or anomalous dispersion, were unsuccessful. The structure was eventually solved by molecular replacement with residues 955-1534 of the AlphaFold structure prediction for MRCK1 (33). The final, experimentally determined, model of MRCK1 955-1534 exhibits a root mean square deviation from the AlphaFold prediction of just 0.906 Å over all atoms (Supplementary Figure S2A). This demonstrates the power of AlphaFold to predict not just protein structures, but intramolecular assemblies of domains from pure co-evolutionary variance (34).

The structure of the C-terminal module of MRCK1 (Figure 2A) reveals an intimate association of the C1, PH and CNH domains. The C1 domain is held together by two zinc ions and makes extensive contacts with both the PH and CNH domains. The PH domain is distinguished from homologous PH domains by a loop insertion between strands β6 and β7 that packs against the CNH domain (Supplementary Figure S2B). The CNH domain forms a 7-bladed beta propeller with strongest homology to the CNH domain of yeast Rho1 (35). The CNH domain is held together by a beta strand contributed by the linker between the PH domain and blade 1 of the beta propeller (Figure 2B, yellow) and the C-terminus of the construct (Figure 2A, dark brown) which together form blade 7. The buried surface area in the interface between the C1-PH and CNH modules of the complex is 1998 Å^2^, with a solvation free energy of −14.9 kcal/mol, indicating that this is a stable association of the three domains. This is also consistent with failed efforts to purify the isolated CNH domain.

**Figure 2.**
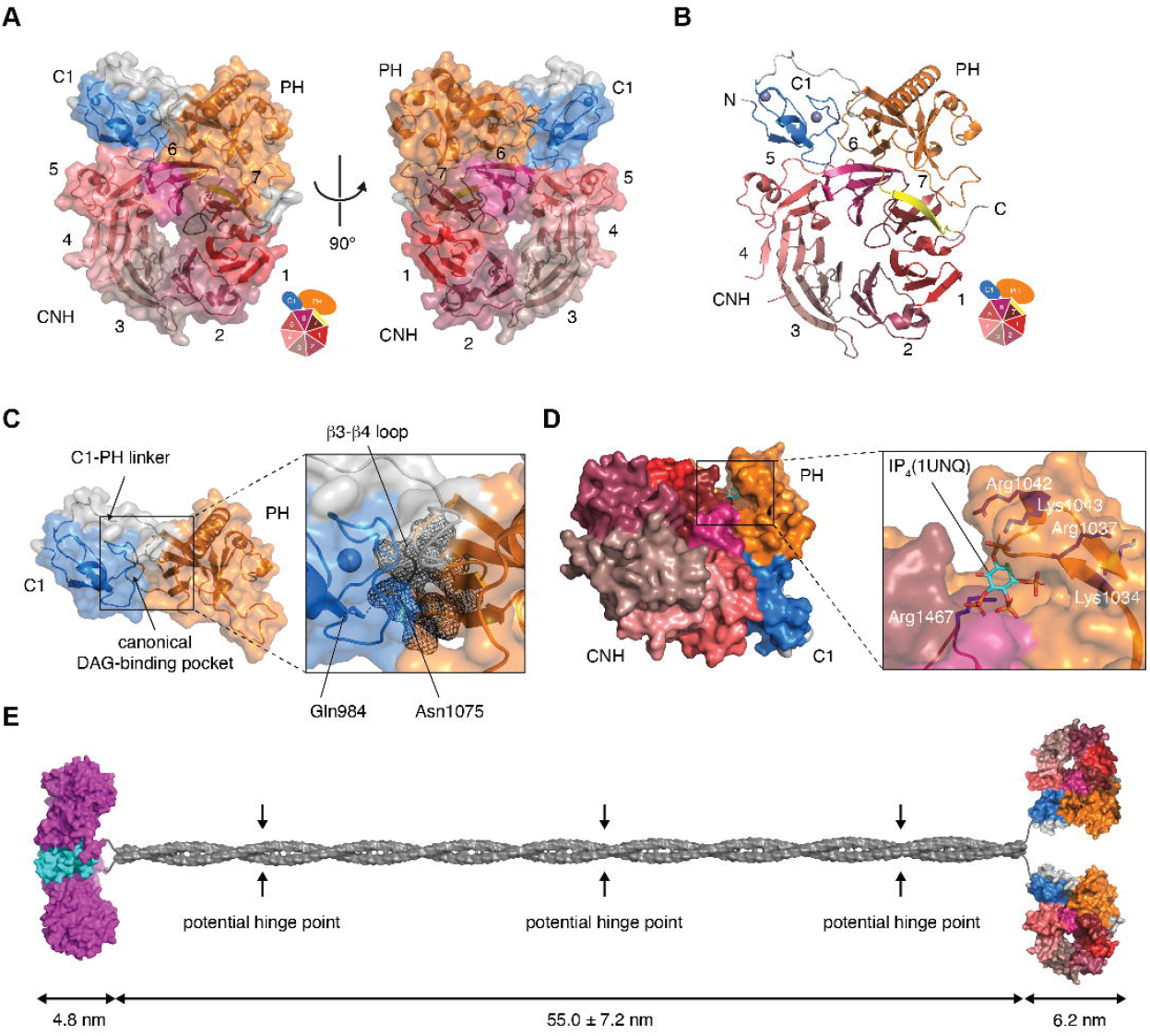
Structure and properties of the C1-PH-CNH module of MRCK1. A. Structure of the C1-PH-CNH module of *C. elegans* MRCK1. (C1, blue; PH, orange; CNH, shades of red). The seven blades of the CNH domain beta propeller are indicated in order. B. Topology of the CNH domain beta propeller. The last beta strand of blade 7 (yellow) is contributed by the linker between the PH domain and blade 1 of the CNH domain. The remaining beta strands of blade 7 are contributed by the C-terminus of the CNH domain. C. Occlusion of the DAG binding pocket of the C1 domain by the β3-β4 loop of the PH domain. D. Superposition of the Akt1 PH domain bound to inositol-1,3,4,5-tetrakisphosphate (IP_4_) with the PH domain of MRCK1. The IP4 ligand is shown in stick representation, the rest of the Akt1 PH domain is omitted for clarity. E. Composite model of MRCK1 based on the coordinates of the kinase domain of MRCKα, a canonical coiled-coil of 55 nm (modeled on the basis of the electron microscopy analysis of MRCK1), and the structure of the C1-PH-CNH module reported in this study. Arrows indicate predicted discontinuities/breaks in the coiled-coil domain that may permit some structural flexibility.

Surface conservation analysis revealed that the C1-PH-CNH module exhibits conservation on the composite surface formed by the C1 and PH domains, but surprisingly little elsewhere (Supplementary Figure S2C top view). Surface electrostatic potential analysis (36) revealed that this surface is predominantly hydrophobic in nature (Supplementary Figure S2D), perhaps indicative of a role in a hereto uncharacterized protein-protein or protein-membrane interaction. Analysis of the structure with PDBePISA (Proteins, Interfaces, Structures and Assemblies) (37, 38) revealed no predicted quaternary assemblies of the C1-PH-CNH module in the crystal lattice.

C1 domains are so called due to their homology to the first conserved domain of protein kinase C (PKC) and are divided into two groups: typical C1 domains that bind the neutral lipid diacylglycerol (DAG) and atypical C1 domains that do not (39). Those C1 domains that bind DAG do so via a cleft formed by unzipped beta strands in which a hydrogen bond network involving conserved glycine and glutamine residues in the cleft is critical for binding the 3’ hydroxyl headgroup of DAG (40). While MRCK and CRIK orthologs all exhibit conservation of this glutamine (Gln984), the putative DAG binding cleft is obscured by a loop between strands β3 and β4 of the PH domain in the structure of MRCK1 (Figure 2C). Within this loop, Asn1075 inserts its side chain into the cleft and makes a hydrogen bond with Gln984. B-factor analysis of the tripartite interface between the C1 domain, the linker between blades 5 and 6 of the CNH domain, and the PH domain indicates a well-ordered interface (Supplementary Figure S2E), although sequence conservation in the β3-β4 loop of the PH domain is poor (Supplementary Figure S2C). This raises obvious questions regarding the conformation of the loop in solution and whether the cleft of the C1 domain is accessible. Although no evidence for DAG binding by the C1 domain of any MRCK protein has yet been presented, binding to phorbol esters, a widely used proxy for DAG, has been reported (41).

Since MRCK has been heavily implicated in the regulation of cytoskeletal processes beneath the plasma membrane and interacts with the membrane-associated small GTPase Cdc42, it is conceivable that its PH domain might be a membrane-interacting domain with specificity for phospholipids. While superposition of the PH domain of Akt1 bound to inositol-1,3,4,5-tetrakisphosphate (IP4) reveals steric clashes with all four phosphate groups of the ligand (Figure 2D), superposition of the PH domain of Evectin-2 bound to phosphatidylserine (PS) reveals fewer clashes. Interestingly, loop residues 1037-1045 between strands β1 and β2 of the MRCK PH domain, which exhibit higher B-factors, show remarkable sequence conservation among MRCK orthologs, including four basic residues (Figure 2D, Supplementary Figure S2F). Blade 7 of the CNH domain contributes an additional basic residue (Arg1467) at the base of the pocket (Figure 2D). While the β1-β2 loop corresponds to residues 11-19 of the Evectin-2 PH domain, which participate in specific interactions with the headgroup of phosphatidylserine (42), none of the basic residues involved in coordinating the phosphate groups are conserved in their 3-dimensional positions. Nevertheless, the presence of a conserved basic loop in the PH domain of MRCK that corresponds to the phospholipid binding pocket of other PH domains is a strong indication that it may participate in membrane binding. Finally, it is worth noting that the β3-β4 loop of the PH domain that inserts Asn1075 into the cleft of the C1 domain forms a surface immediately adjacent to the putative phospholipid binding β1-β2 loop. As such, binding of a ligand to the C1 domain would necessarily displace the β3-β4 loop of the PH domain and potentially impact the properties of the phospholipid binding pocket.

In summary, the C1-PH-CNH module of MRCK1 forms a globular structure in which the putative membrane binding surfaces of the C1 and PH domains are coaligned. The role of the CNH domain is not clear from the structure, but presumably facilitates additional protein-protein interactions or protein-membrane interactions that localize MRCK activity to the appropriate subcellular compartment. By combining the electron microscopy images of full-length MRCK1 (Figure 1D) and the high-resolution structure of the regulatory domains (Figure 2A) with the structure of the dimeric kinase domains of human MRCKα (PDB 4AW2, unpublished), we propose a composite high-resolution model of MRCK1 in which the kinase and regulatory domains are linked by a coiled-coil of 55 nm (Figure 2E). While the model depicts a continuous coiled-coil and the electron micrographs depict a semi-rigid, extended coiled-coil domain, predicted discontinuities in the coiled-coil (Supplementary Figure 1A) may introduce ‘hinges’ that could permit some flexibility.

### Membrane-binding properties of the C1-PH-CNH module of MRCK1

To assess if MRCK1 can directly interact with membranes, we first determined the binding capacity of MRCK1 to liposomes reconstituted from a Folch fraction of brain lipids by liposome pelleting assay. These liposomes were capable of binding approximately 50% of full-length MRCK1 or MRCK1^955-1534^ at a total lipid concentration of 1 mM and 1.5 μM MRCK1 (Figure 3A). The interaction was specific for the liposomes, as MRCK1 remained entirely in the supernatant in their absence. To investigate which lipids could be specifically recognized by MRCK1 at the membrane surface, we probed the interaction of its regulatory module C1-PH-CNH to membrane surrogates containing various combinations of lipids in a systematic manner using the high-throughput liposome microarray assay (LiMA) (43, 44). In parallel, we probed the binding of the PH domain of phospholipase C (PLC) δ as a control of membrane binding specificity (45). Membrane lipid binding was quantified by determining the ratio of fluorescent protein to that of Atto647-conjugated phosphatidylethanolamine (Atto647-PE) contained in liposomes. In this context, we observed that MRCK1^955-1534^-EGFP bound preferentially to bis- and tris-phosphorylated phosphoinositides (PIPs) over mono-phosphorylated PIPs in a dose dependent manner, as well as to C18 ceramide-1-phosphate but to a lesser extent (Figure 3B, Supplementary Figure S3A and S3B).

**Figure 3.**
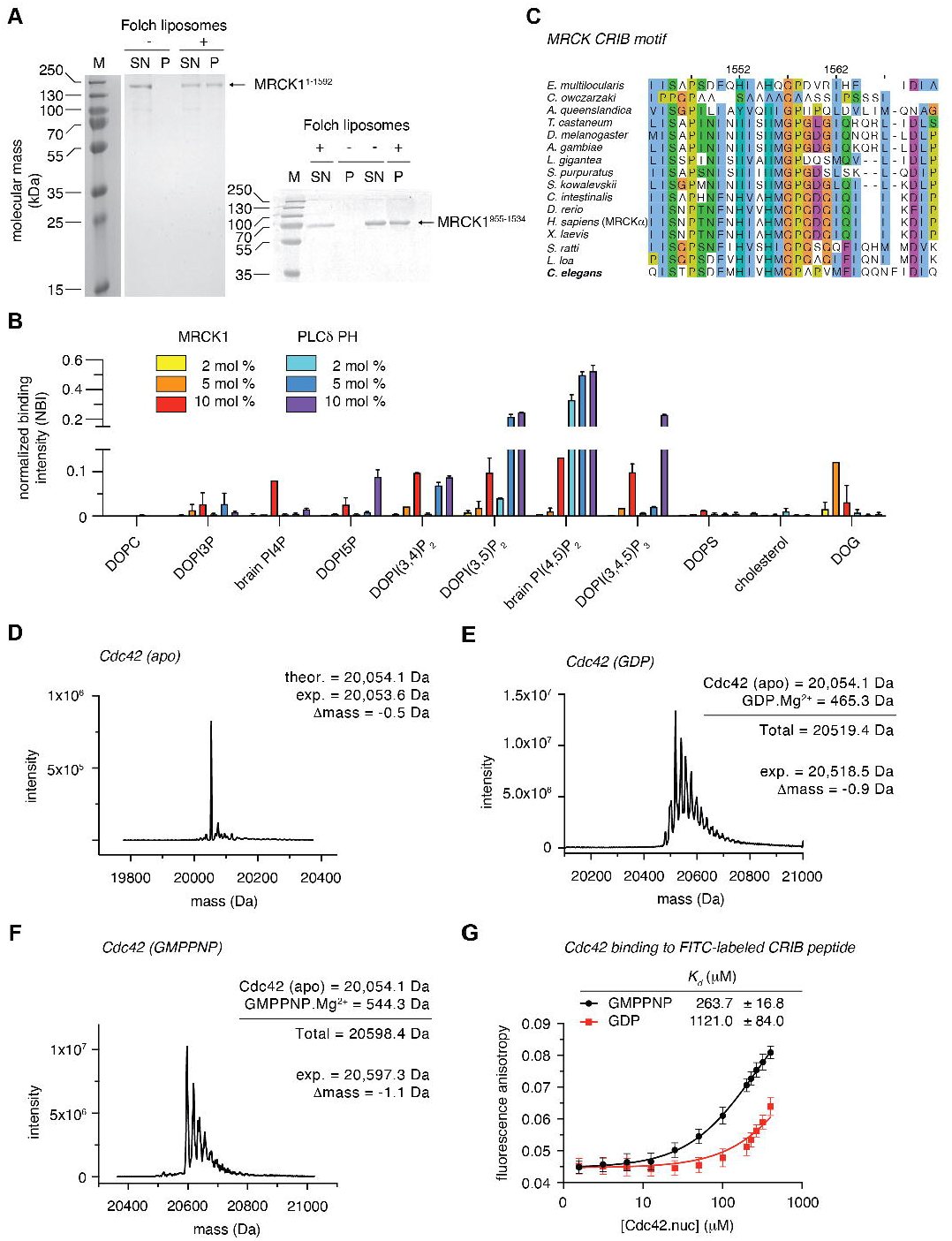
Membrane-binding properties of the C1-PH-CNH module of MRCK1. A. Binding of MRCK1 to Folch liposomes. Full-length MRCK1 (MRCK1^1-1592^) as well as the C1-PH-CNH module (MRCK1^955-1534^) exhibited approximately 50% binding to Folch liposomes at 1 mM total lipid concentration. B. Normalized binding of MRCK1^955-1534^-EGFP (C1-PH-CNH) to liposomes containing the indicated lipid species in a high-throughput LiMA assay. Binding was quantified as the ratio of MRCK1^955-1534^-EGFP to Atto647-conjugated phosphatidylethanolamine incorporated into the liposomes at a fixed concentration of x. MRCK1 binding to each lipid was evaluated with three different concentrations of signaling lipid: yellow (2 mol %), orange (5 mol %), red (10 mol %) and compared to the PH domain of PLCδ, which binds specifically to PI(4,5)P2 and was used as a positive control: cyan (2 mol %), blue (5 mol %), purple (10 mol %). C. Sequence alignment of the CRIB domain of MRCK orthologs. The CRIB domain of *C. elegans* MRCK1 is highlighted in bold. D. Intact, denaturing mass spectrometry of purified, C-terminally truncated Cdc42 (residues 2-178). E. Native mass spectrometry of GDP-loaded Cdc42. F. Native mass spectrometry of GMPPNP-loaded Cdc42. G. Binding of Cdc42 to a FITC-labeled CRIB peptide corresponding to the *C. elegans* CRIB motif sequence shown in panel C, determined by fluorescence anisotropy. Data fitted with a one site binding model to determine the affinity of interaction.

*In vivo,* the specific targeting of MRCK1 to apical membranes may be explained by two additional factors: while MRCK1^955-1534^ is monomeric in solution, full-length MRCK1 is a dimer, which doubles the membrane interacting surface; secondly, MRCK1, like all MRCK orthologs, encodes a Cdc42 and Rac-interactive binding (CRIB) domain C-terminal to its CNH domain (Figure 1A and 3C). It is therefore likely that membrane-associated MRCK is tethered via a set of polyvalent interactions, including the binding to two copies of membrane-anchored Cdc42. To determine the affinity of MRCK1 for Cdc42, we expressed a C-terminally truncated Cdc42 lacking its geranylgeranylation motif in *E. coli* and purified it to homogeneity. The mass of the recombinant protein was confirmed by intact, denaturing mass spectrometry (Figure 3D). Using the same protocol as we previously reported for RhoA (7), we loaded Cdc42 with either guanosine 5’-diphosphate (GDP) or the non-hydrolyzable GTP analog guanosine 5’-[(β,γ)-imido]triphosphate (GMP-PNP). Stoichiometric loading of Cdc42 with either GDP or GMP-PNP was confirmed by native mass spectrometry (Figure 3E and Figure 3F, respectively).

We measured the binding affinity of Cdc42 to a peptide comprising the CRIB motif of MRCK1 (residues 1543-1572) by fluorescence anisotropy. While Cdc42 bound to GDP exhibited an estimated *K_d_* in the millimolar range, the binding to Cdc42 loaded with GMP-PNP was 5-fold stronger (Figure 3G). The binding of MRCK1 to two molecules of membrane-anchored Cdc42 is therefore likely to be in the low-mid nanomolar range; if additional specific interactions with lipids are factored in, this will be even tighter.

## Discussion

We have determined that full-length MRCK1 adopts a highly extended conformation *in vitro,* with dimeric kinase domains at its N-terminus and monomeric membrane- and Cdc42-binding domains at its C-terminus, joined by a coiled-coil of approximately 55 nm in length. MRCK1 binds specifically to active, GTP-bound Cdc42 via its CRIB motif and weakly to membranes, with a preference for di-phosphorylated phosphoinositides. While the structural basis for these protein-lipid interactions will require further investigation, the combination of membrane-anchoring regulatory domains at one end of a highly extended particle with kinase domains at the other suggests that the coiled-coil domain of MRCK, like in ROCK, may act as a molecular ruler to position its kinase domains at a fixed distance from a membrane scaffold.

The defining members of the DMPK sub-family of AGC kinases (DMPK, CRIK, MRCK, and ROCK) exhibit a conserved domain architecture with the principal difference being the length of their coiled-coil domains. The coiled-coil domains of each family member are evolutionarily conserved in length, much less in sequence. Each member has been reported to associate with the plasma membrane either directly, via their C-terminal regulatory domains or a transmembrane helix, or indirectly via binding to a membrane-anchored small GTPase. Structures of the kinase domains of ROCK1, ROCK2, MRCKα (PDB 4AW2, unpublished), MRCKβ and DMPK1 all reveal a dimeric, head-to-head conformation mediated by the CHB domains, which clamp the C-terminal extension of the kinase domains, thereby maintaining an active conformation (27–29). Deletion or mutation of the hydrophobic motif in the C-terminal tail of ROCK leads to abrogation of dimerization and a corresponding loss of kinase activity (46). The unphosphorylated activation loops of these AGC kinases adopt a stable conformation compatible with substrate binding and each has been reported to exhibit robust kinase activity *in vitro.* MRCK, DMPK, and CRIK do not possess a phosphorylatable residue at the canonical position in their activation loops, while substitution of the activation loop threonine for alanine or a phosphomimetic glutamate in ROCK does not change its specific activity (7). Allosteric regulation of kinase activity by phosphorylation, protein co-factors, membrane-binding or intramolecular autoinhibition has either not been reported or early reports, particularly in the case of ROCK (30, 47–49), have not been further substantiated with purified recombinant full-length protein (7). While an early report claimed autoinhibition of MRCKα activity by a distal region of the coiled-coil domain and modest (2-fold) activation by phorbol esters (50), we could not detect any evidence for a physical association between the C-terminal regulatory domains and the kinase domain in full-length MRCK1. Previous work has also shown that overexpression of active Cdc42 did not influence the activity of immunoprecipitated MRCK (9), which is consistent with the C-terminal location of the flexibly linked CRIB motif, ~60 nm away from the kinase domains.

Taken together, these observations hint at other, cellular mechanisms, that likely regulate signaling via these kinases. One such mechanism, which we have previously proposed for ROCK, is the concept of a molecular ruler, in which the length of the coiled-coil domain fixes the kinase domains at a precise molecular location with respect to the membrane. Truncation mutants, in which the coiled-coil of ROCK was shortened while preserving catalytic activity, elicited a defect in stress fiber formation when expressed in cells, indicating that the length of the coiled-coil is important for ROCK function (7). Such a mechanism necessarily relies on the precise positioning of enzyme and substrate within a non-diffusive framework, which could be argued to be the case in the context of the regulation of actomyosin contractility beneath the plasma membrane.

Within the actin cortex, which microscopic studies have measured to span a distance of approximately 20-200 nm beneath the plasma membrane in various mammalian cells (51), actomyosin fibers are precisely arranged in order to effect changes in cell shape and motility. MLC2, phosphorylation of which regulates this contractility, is bound to myosin II, while MLCP must also be recruited to MLC2 in order to affect its dephosphorylation. Experimental characterization of DMPK, MRCK, and ROCK, together with bioinformatic analysis of CRIK, suggests that the four DMPK family members could, in principle, span distances from as short as 15 nm to as far as 145 nm from the plasma membrane (Figure 4), thereby differentially controlling actomyosin contractility within the actin cortex. The integration of multiple membrane-based signals, including protein-lipid and protein-protein interactions, likely specify the target membrane to which these kinases are localized. Both ROCK and MRCK exhibit subcellular localization to the apical plasma membrane (52, 53), where they play roles in the establishment of epithelial cell polarity and apical constriction. The specificity of signaling is further orchestrated by a variety of adaptor proteins that physically link ROCK, MRCK, and CRIK to components of the actin cytoskeleton. These adaptors include Shroom, an actin-binding protein that binds to the coiled-coil domain of ROCK via its SD2 domain, and has been shown to be essential for apical constriction (23, 54, 55), and leucine repeat adaptor proteins 25 (LRAP25) and 35a (LRAP35a), which link MRCK to LIM-kinase (LIMK) (56) and MYO18A (57) respectively. While the interaction site of LRAP35a was mapped to a so-called kinase-inhibitory motif (KIM) in the coiled-coil domain of MRCKα (57) previously determined to be an autoinhibitory domain (50, 58), we observed no evidence of a physical association of the coiled-coil domain with either end of MRCK1 in the electron micrographs.

**Figure 4.**
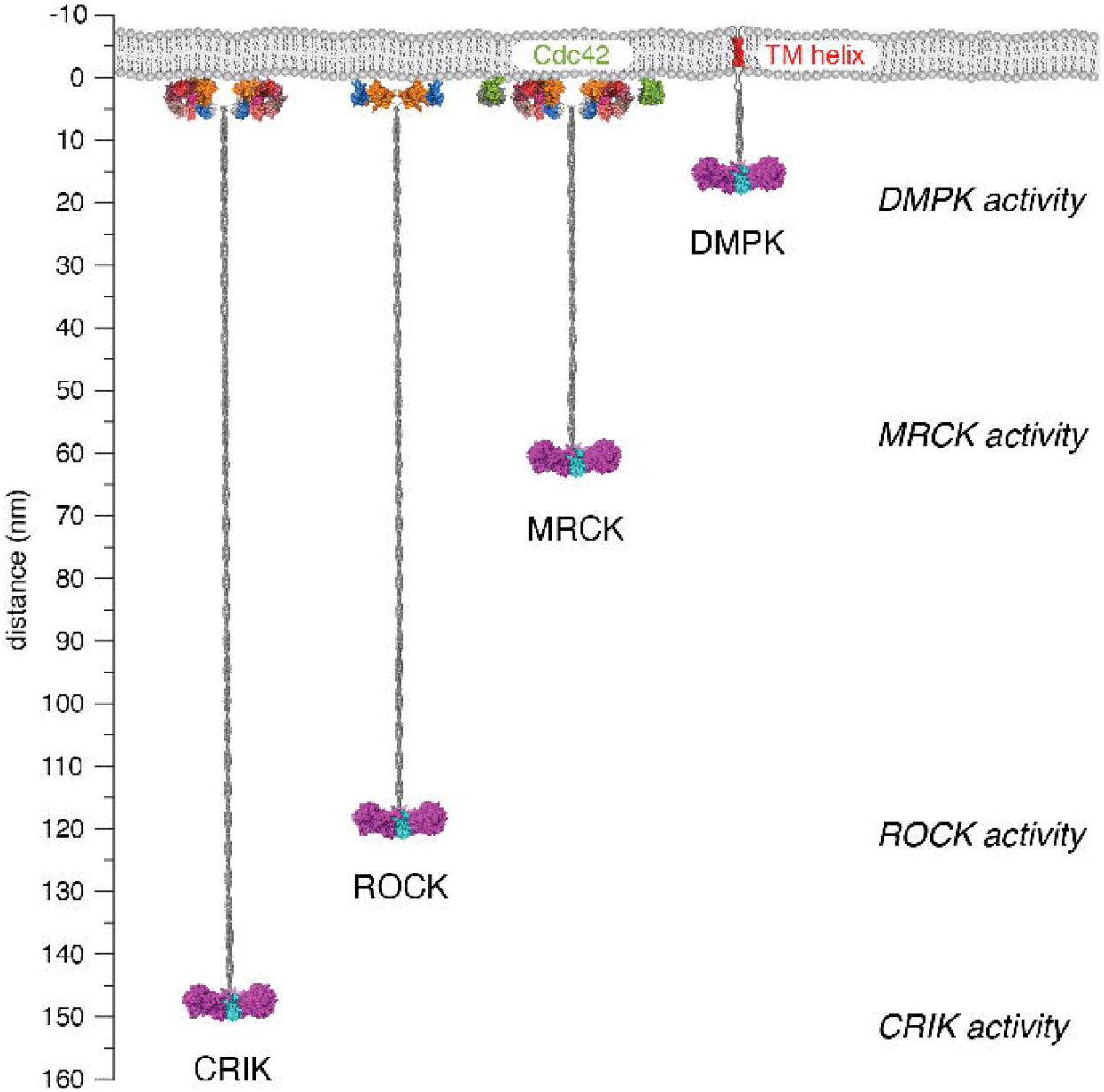
Model of DMPK kinases as molecular rulers. The four DMPK family members (CRIK, DMPK, MRCK and ROCK) are depicted bound to a hypothetical membrane. Specific membrane anchors are indicated where known. The coiled-coil domains are depicted at their maximum extensions based on experimental analysis of DMPK, MRCK1 and ROCK2 together with bioinformatic analysis of all four family members. A ruler indicates the distance that they are capable of spanning between the membrane and a hypothetical position in the actin cortex.

The membrane localization of all members of the DMPK family is by now quite well established, though there is much still to learn about the specificity of the interactions that control their precise subcellular localization. Over-expression of the regulatory domains of ROCK and MRCK is sufficient to induce the dominant negative inhibition of stress fiber formation (48, 59) and neurite outgrowth (13), presumably by the displacement of the endogenous kinase. In the case of MRCK, ectopic expression of a regulatory domain construct lacking its CRIB motif is sufficient to impair neurite outgrowth (13), indicating that, in addition to its recruitment by Cdc42, specific membrane interactions most likely contribute to its localization. However, while binding of the C1 domain of MRCKα to phorbol esters has previously been described (41, 50), we did not detect robust, concentration-dependent binding to its putative physiological ligand DAG in our LiMA assay (Figure 3A). We observed incomplete binding of both full-length MRCK1 and its isolated regulatory domains to Folch liposomes *in vitro,* which may be a consequence of incorrect membrane topology (high positive curvature), lipid composition, or a combination thereof. The binding of the regulatory domains to di-phosphorylated phosphoinositides will need further investigation, but conservation of the putative phospholipid binding pocket suggests that specific lipid binding may be functionally relevant.

Although full-length CRIK and DMPK1 have, as yet, not been characterized, they also exhibit conservation of the length of their coiled coils. DMPK orthologs are found only in tetrapods, where they have been implicated primarily in muscle-specific functions. Bioinformatic analysis reveals that their predicted coiled-coil is invariant in length (8.9 nm), while the linker length between the N-terminal kinase domain and the coiled-coil domain varies between 49 and 118 aa (Supplementary Figure S4A). Approximately 75 amino acids that lie between the coiled-coil domain and the transmembrane helix are conserved in both sequence and length, but have no known function. The evolution of deuterostomes coincides with a remarkable conservation of the length of CRIK, with a predicted coiled-coil domain of 145.9 ± 1.4 nm (Supplementary Figure S4B-D). CRIK orthologs are found in some, but not all protostomes, where they exhibit more variable length in their coiled-coil domain. DMPK1 contains an N-terminal kinase domain with the same CHB domain-mediated dimeric arrangement as found in ROCK and MRCK (29) followed by a short coiled-coil and a C-terminal transmembrane helix. While X-ray crystallographic analysis of the isolated coiled-coil domain of DMPK has revealed a trimeric arrangement, constructs encompassing both the kinase domains and the coiled-coil domain are exclusively dimeric in solution (60).

In summary, our findings are compatible with a model in which the activity of DMPK kinases is governed by the precise positioning of their kinase domains with respect to the membrane scaffold (Figure 4). This, we argue, may be accomplished by a ‘molecular ruler’ in the form of the coiled-coil domain. While the model depicted is clearly an oversimplification of the cellular architecture of the membrane and the actin cortex beneath the membrane, it provides a framework for future work that is required to understand why the length of these coiled-coil domains have been so strongly conserved during evolution and which molecular and cellular events are required to bring enzyme (kinase) and substrate into proximity, as well as the correct orientation required for catalysis (61).

## Acknowledgements

We would like to thank Montserrat Soler Lopez for assistance in data collection on ID29 at the European Synchrotron Radiation Facility (ESRF), Grenoble, France. We would also like to thank Dorothea Anrather and Markus Hartl in the Max Perutz Labs Mass Spectrometry Facility for help with the acquisition of mass spectra using instruments of the Vienna BioCenter Core Facilities (VBCF). We are grateful to Marlene Brandstetter in the VBCF Electron Microscopy Facility for help with preparing rotary-shadowed samples of MRCK1. This work was supported by a Hertha Firnberg Postdoctoral Fellowship to L.T. and FWF grants P28135, P30584, and P33066 to T.A.L.

C.G. was supported by the EMBL Interdisciplinary Postdoc Programme under Marie Curie COFUND Actions. A.C.G. acknowledges the financial support of the Louis-Jeantet Foundation.

## Author contributions

L.T. purified all recombinant proteins and performed all the biochemical experiments reported in this study. E.F. performed liposome pelleting assays with MRCK1 *in vitro.* L.T. performed LiMA assays with the help of C.G. in the lab of A.C.G. T.A.L. collected diffraction data on MRCK1 crystals grown by L.T. and solved the structure. L.T. built and refined the structure. L.T., A.C.G. and T.A.L. obtained the funding to support the work.

## Materials and Methods

### Protein expression and purification

MRCK1^1-1592^ (full length protein) and MRCK^955-1534^ were expressed in Sf9 insect cells with a N-terminal His_10_-StrepII tag. For MRCK1 full length purification cells were lysed in 30 ml lysis buffer (50 mM Tris pH 7.5, 140 mM KCl, 10 mM NaCl, 10 mM ß-glycerophosphate, 2 mM sodium pyrophosphate, 1 mM TCEP, 2 mM Benzamidine, 0.2 mM sodium orthovanadate, 40 mM NAF, 1 mM PMSF, 100 μM Bestatin, 14 μM E-64,10 μM Pepstatin and 1 μM Phosphoramidon). After one freeze/thaw cycle, 1 mM MgCl_2_ and Benzonase were added and samples were left on ice for 30 min. After centrifugation @ 18000 rpm for 30 min @ 4°C samples were loaded to a His Trap FF column, which was equilibrated in buffer A (50 mM Tris pH 7.5, 140 mM KCl,1 mM TCEP, 50 mM NAF). Column was washed with 2% buffer B (50 mM Tris pH 7.5, 140 mM KCl, 1 mM TCEP, 50 mM NAF, 1 M Imidazole) and 4% buffer B before gradient elution into 100% buffer B. Fraction were collected in tubes containing 50 mM Tris pH 7.5, 100 mM KCL, 1 mM TCEP to reduce the imidazole concentration by two third. Peak fractions were pooled, diluted 1:3 in QA buffer (50 mM Tris pH 8.0, 1 mM TCEP) and TEV cleaved over night @ 4°C. The next day sample was loaded to a HiTrap Q column equilibrated in QA buffer and eluted with a gradient into 100 % buffer QB (50 mM Tris pH 8.0, 1 mM TCEP, 1 M NaCl). Elution fractions were pooled and concentrated and buffer exchanged using an amicon concentrator (30 kDa cutoff) into storage buffer (20 mM Tris pH 8.0,1 mM TCEP, 100 mM KCl and frozen @ −80°C.

For purification of MRCK^955-1534^ Sf9 cell pellets were resuspended in lysis buffer (50 mM HEPES pH 8.0, 150 mM KCl, 1 mM TCEP) with 2 mM MgCl2, 2 mM benzamidine, 50 μl 100x protease inhibitor mix (P8849 Sigma), 1 μl benzonase, and 1 mM PMSF. The resuspended cells were incubated on ice for 30 minutes, and then centrifuged for 30 minutes at 18000 rpm, 4°C. The supernatant was loaded onto a 5 ml HisTrap FF column (GE Healthcare Life Sciences) washed and equilibrated in buffer A (50 mM Hepes pH 8.0, 150 mM KCl, 1 mM TCEP). After washing the column with 2% buffer B (buffer A, 1 M imidazole) protein was eluted with a gradient into 100% buffer B. Peak fractions were pooled and 1:1 diluted with lysis buffer. 150 μl of TEV (2 mg/ml) were added and the cleavage reaction was incubated at 4°C over night. The sample was further diluted 1:4 with SA buffer (50 mM Tris pH 8.0, 1 mM TCEP) to lower the salt concentration for binding to a cation exchange column. The sample was then loaded onto a 5 ml HiTrap SP FF column (GE Healthcare Life Sciences) followed by gradient elution with elution buffer SB (50 mM Tris pH 8, 1 M NaCl, 1 mM TCEP). Peak fractions were pooled, concentrated and loaded to a HiLoad 10/300 Superdex 200 column (GE Healthcare Life Sciences) equilibrated with gel filtration buffer (20 mM Tris pH 7.5, 150 mM KCl, 1 mM TCEP, 1% glycerol). Peak fractions were pooled and concentrated to 4 mg ≥ ml and the protein was frozen in LN and stored at −80°C.

Cdc42 was expressed with an N-terminal GST-tag in *E.Coli* over night at 22°C. Cells were harvested and pellets resuspended in 50 ml lysis buffer (50 mM Tris pH 7.4, 150 mM KCL, 2 mM MgCl2, 1 mM TCEP, 2 mM Benzamidine) and frozen at −80°C. After thawing 10 mg Lysozyme and 2 μl Benzonase were added and solution was incubated 30 min on ice. After sonication and centrifugation for 15 min @ 18000 rpm at 4°C supernatant was incubated with 1 ml in lysisbuffer equilibrated GST beads for 3 h @ 4°C rotating. Beads were collected @ 500 g for 10 min and washed with buffer A (50 mM Tris pH 7.4, 150 mM KCL, 2 mM MgCl2, 1 mM TCEP) and beads resuspended in 15 ml buffer A. The protein was incubated with TEV protease over night @ 4°C and subjected to a SD75 16/600 equilibrated in buffer A. Elution fractions were collected, concentrated, frozen in LN and stored @ −80°C.

### Rotary shadowing electron microscopy

Preparation of rotary-shadowed MRCK1 and ROCK2 was carried out as previously described for ROCK2 (7). Samples were diluted to a final concentration of 50 μg ml^-1^ in spraying buffer containing 100mM ammonium acetate and 30% (v/v) glycerol, pH 7.4. After dilution, the samples were sprayed on freshly cleaved mica chips and immediately transferred into a BAL-TEC MED020 high-vacuum evaporator equipped with electron guns. While rotating, the samples were coated with 0.6-nm platinum at an angle of 4–5°, followed by 10-nm carbon at 90°. The obtained replicas were floated off from the mica chips, picked up on 400 mesh Cu/Pd grids and inspected in an FEI T20 G2 transmission electron microscope operated at 80 kV. Electron micrographs were acquired using an FEI Eagle 4k charge-coupled device camera.

### Kinase assay

Kinase assays were done using the ADP-glo kit from Promega according to the manufacturers instructions. Assays were done in white 96 well plates using substrate concentrations between 2 and 24 μM RMLC2. Kinase assays were done for 40 min @ RT in a final volume of 25 μl containing 100 nM full length MRCK1, substrate and 1 mM ATP in 50 mM Tris pH 5.7, 100 mM KCl, 1 mM TCEP, 20 mM MgCl_2_. 25 μl ADP-glo reagent was added afterwards and samples were incubated for another 40 min @ RT to deplete any remaining ATP. For luminescence detection 50 μl kinase detection reagent was added to each sample and incubation was done for 1 hour @ RT. Samples were measured for luminescence in a TECAN Infinite F500 plate reader.

### Crystallization and structure determination

MRCK1^955-1534^ was crystallized in 100 mM Bicine/Tris (base) at pH 8.5, 30 μM of each sodium nitrate, disodium hydrogen phosphate and ammonium sulfate, 20% PEG550 MME, 8% PEG20K, 3% glycerol. Crystals grew in spacegroup P1 with unit cell dimensions a=69.14 Å, b=97.47 Å, c=131.41 Å, α=81.38° β=77.77° γ=80.39°. Crystals diffracted to 2.14 Å resolution and contained 5 molecules per asymmetric unit. The structure was solved by molecular replacement with the AlphaFold prediction for residues 955-1534 of *C. elegans* MRCK1 (AF-O01583-F1). The model was rebuilt with Phaser Autobuild (62), manually checked and regions of ambiguous electron density built in Coot. The final model was refined to *R_free_* = 0.244 and *R_work_* =0.195 (for dataset and model statistics, see Supplementary Table 1).

### Liposome microarray assay (LiMA)

LiMA (43) was performed in the lab of Anne-Claude Gavin (EMBL Heidelberg, Germany) according to (44). MRCK^955-1534^ C-terminally tagged with EGFP and a positive control, PLCd1-PH fused to sfGFP, were applied to microarrays printed with different signaling lipids. In brief, lipids of interest were combined with the carrier lipid DOPC, PEGylated PE and PE labelled with Atto 647 dye (PE-Atto 647, 0.1 mol%). Such lipid mixtures contained 2, 5 and 10 mol% of the signaling lipid and were spotted onto a thin agarose layer (TAL). The agarose layers were hydrated using Buffer A: 20 mM Tris-HCl pH 8.5, 200 mM NaCl and vesicles formed spontaneously. Efficiency of liposome formation was verified by fluorescence microscopy. The protein was diluted to 7μM in buffer A and 40 μl were subjected to each array. Microarrays were incubated for 20 min at RT and subsequently washed three times with 40 μl of buffer A. Chips were analyses by automated fluorescence microscopy. Positions of liposomes were determined by tracking the fluorescence of PE-Atto 647 and pictures were taken at 3 ms and 5 ms. In parallel the fluorescence of EGFP was determined at eight different exposure times, namely at 1, 5, 10, 30, 75, 100, 200 and 300 ms. Pictures were processed using CellProfiler and CPAnalyst. Only EGFP signals that overlapped with Atto 647 signals were taken into account. Normalized binding intensity (NBI) was calculated as a ratio between EGFP and Atto 647 fluorescence, normalized by exposure time. Three microarrays were examined, carrying liposomes with the following signaling lipids; PIP-chip: DOPA, DOPE, DOPI, DOPS, DODAG, cardiolipin, BMP, DOPI(4,5)P2, DOPG; GLP-chip: Ceramide C16, ceramide(1)P C16, ceramide(1)P C18, S(1)P, S, SM, DOPI(4,5)P2, DOPS; SL-chip: DOPI(3)P, brain PI(4)P, DOPI(5)P, DOPI(3,4)P2, DOPI(3,5)P2, brain PI(4,5)P2, DOPI(3,4,5)P3, DOPS and cholesterol. Each microarray was done in triplicates.

### Rhodamine-labeled sucrose-loaded vesicles (SLVs)

Folch fraction I lipids (25 mg / ml in chloroform) were pipetted into borosilicate test tubes (1 ml of 1 mM solution gives in the end 20 sample runs) and 0.006 mM Rhodamine-DHPE was included. Lipids were dried to a thin film using a dry nitrogen stream. 1 ml of 20 mM HEPES, pH 7.5, 0.3 M Sucrose was added and samples were vortexed vigorously. Samples were split into 4 x 250 μl aliquots and 4 freeze-thaw cycles in liquid nitrogen followed by thawing in sonicating water bath were done. To 250 μl SLVs, 300 μl 20 mM HEPES, pH 7.5, 100 mM KCl were added and samples were spinned @ 80000 rpm for 30 min @ RT in a TLA 100.1 rotor. The supernatant was removed and pellets were resuspended in 250 μl 20 mM HEPES, pH 7.5 and 100 mM KCl.

### Liposome pelleting assay

For liposome pelleting assays 1.5 μM full length MRCK1 or MRCK^955-1534^ were incubated with or without liposomes at RT for 30 min. Final concentration of liposomes was 0.5 mg / ml. Samples were centrifuged in an ultracentrifuge at 21000 rpm using a TLA 100 rotor for 30 min at RT. Supernatant and pellet fractions were evaluated by densitometry of Coomassie-stained SDS-PAGE gels.

### Cdc42 nucleotide exchange

Cdc42^2-178^ (co-purified with a mixture of GDP and GTP) was incubated with Dbs exchange factor (5:1) in a 100-fold molar excess of GMPPNP, 30 min at RT. Cdc42^2-178^ was separated from Dbs and free nucleotide by size exclusion chromatography. The extent of GMPPNP loading was determined by native mass spectrometry (see below).

### Mass spectrometry

The protein mass was determined on a Synapt G2-Si Q-ToF mass spectrometer (Waters) coupled to an Ultimate 3000 HPLC system (Dionex, Thermo Fisher Scientific) via a ZSpray ESI source (Waters). Proteins were separated on an Aeris Widepore C4 column (3.6 μm particle size, dimensions 2.1 x 150 mm, Phenomenex) with a working temperature of 55 °C, applying a 10 min gradient from 8 % to 63 % acetonitrile in 0.1 % formic acid at a flow rate of 300 μL/min. On the mass spectrometer the sampling cone voltage was 40 V, the desolvation gas temperature 450 °C and the source temperature 120 °C.

For MS experiments under native conditions, the buffer of the protein solutions was exchanged to 50 mM ammonium acetate, pH 6.8. Measurements were performed on the Synapt G2-Si equipped with the NanoLockSpray electrospray source (Waters) using pre-opened PicoTip emitters (New Objective). Electrospray voltage was set at 1.8 kV, the source temperature at 80 °C and sampling cone voltage at 80 V. All data were analysed in MassLynx V 4.1 using the MaxEnt 1 process to reconstruct the uncharged average protein mass.

### Fluorescence anisotropy

The binding affinity of the C. elegans MRCK1 CRIB motif (residues 1543-1572) was determined by titration of purified Cdc42^2-178^ loaded either with GDP or GMPPNP into a buffer containing 100 nM fluorescein-labeled peptide. Fluorescence anisotropy was measured on a Perkin-Elmer LS50 fluorimeter with λ_ex_ = 500 nm and λ_em_ = 518 nm, at 20 °C in 20 mM Tris, pH 8.0, 100 mM NaCl, and 1 mM TCEP. Each concentration of Cdc42^2-178^ was measured 50 times with an integration time of 1 s and the mean plotted. The error bars represent the SD of the measurements. Three independent titrations were performed. The obtained binding curves were fit with a one-site binding model to estimate the binding affinity.

## References

1. Svitkina T (2018) The Actin Cytoskeleton and Actin-Based Motility. Cold Spring Harb Perspect Biol 10(1):a018267.

2. Adelstein RS, Anne Conti M (1975) Phosphorylation of platelet myosin increases actin-activated myosin ATPase activity. Nature 256(5518):597–598.

3. Pires EM, Perry S V (1977) Purification and properties of myosin light-chain kinase from fast skeletal muscle. Biochem J 167(1):137–46.

4. Shimizu H, Ito M, Miyahara M, Ichikawa K, Okubo S, Konishi T, Naka M, Tanaka T, Hirano K, Hartshorne DJ (1994) Characterization of the myosin-binding subunit of smooth muscle myosin phosphatase. J Biol Chem 269(48):30407–11.

5. Shirazi A, Iizuka K, Fadden P, Mosse C, Somlyo AP, Somlyo A V, Haystead TA (1994) Purification and characterization of the mammalian myosin light chain phosphatase holoenzyme. The differential effects of the holoenzyme and its subunits on smooth muscle. J Biol Chem 269(50):31598–606.

6. Alessi D, MacDougall LK, Sola MM, Ikebe M, Cohen P (1992) The control of protein phosphatase-1 by targetting subunits. The major myosin phosphatase in avian smooth muscle is a novel form of protein phosphatase-1. Eur J Biochem 210(3):1023–35.

7. Truebestein L, Elsner DJ, Fuchs E, Leonard TA (2015) A molecular ruler regulates cytoskeletal remodelling by the Rho kinases. Nat Commun 6:1–13.

8. Amano M, Ito M, Kimura K, Fukata Y, Chihara K, Nakano T, Matsuura Y, Kaibuchi K (1996) Phosphorylation and Activation of Myosin by Rho-associated Kinase (Rho-kinase). J Biol Chem 271(34):20246–20249.

9. Leung T, Chen X-Q, Tan I, Manser E, Lim L (1998) Myotonic Dystrophy Kinase-Related Cdc42-Binding Kinase Acts as a Cdc42 Effector in Promoting Cytoskeletal Reorganization. Mol Cell Biol 18(1):130–140.

10. Wilkinson S, Paterson HF, Marshall CJ (2005) Cdc42-MRCK and Rho-ROCK signalling cooperate in myosin phosphorylation and cell invasion. Nat Cell Biol 7(3):255–261.

11. Kimura K, Ito M, Amano M, Chihara K, Fukata Y, Nakafuku M, Yamamori B, Feng J, Nakano T, Okawa K, Iwamatsu A, Kaibuchi K (1996) Regulation of Myosin Phosphatase by Rho and Rho-Associated Kinase (Rho-Kinase). Science (80-) 273(5272):245–248.

12. Tan I, Ng CH, Lim L, Leung T (2001) Phosphorylation of a novel myosin binding subunit of protein phosphatase 1 reveals a conserved mechanism in the regulation of actin cytoskeleton. J Biol Chem 276(24):2I209–I6.

13. Chen X-Q, Tan I, Leung T, Lim L (1999) The Myotonic Dystrophy Kinase-related Cdc42-binding Kinase Is Involved in the Regulation of Neurite Outgrowth in PC12 Cells. J Biol Chem 274(28):19901–19905.

14. Madaule P, Eda M, Watanabe N, Fujisawa K, Matsuoka T, Bito H, Ishizaki T, Narumiya S (1998) Role of citron kinase as a target of the small GTPase Rho in cytokinesis. Nature 394(6692):491–494.

15. Marston DJ, Higgins CD, Peters KA, Cupp TD, Dickinson DJ, Pani AM, Moore RP, Cox AH, Kiehart DP, Goldstein B (2016) MRCK-1 Drives Apical Constriction in C. elegans by Linking Developmental Patterning to Force Generation. Curr Biol 26(16):2079–2089.

16. Gally C, Wissler F, Zahreddine H, Quintin S, Landmann F, Labouesse M (2009) Myosin II regulation during *C. elegans* embryonic elongation:LET-502/ROCK, MRCK-1 and PAK-1, three kinases with different roles. Development 136(18):3109–3119.

17. Gomes ER, Jani S, Gundersen GG (2005) Nuclear movement regulated by Cdc42, MRCK, myosin, and actin flow establishes MTOC polarization in migrating cells. Cell 121(3):451–63.

18. Winter CG, Wang B, Ballew A, Royou A, Karess R, Axelrod JD, Luo L (2001) Drosophila Rho-associated kinase (Drok) links Frizzled-mediated planar cell polarity signaling to the actin cytoskeleton. Cell 105(1):81–91.

19. Zihni C, Vlassaks E, Terry S, Carlton J, Leung TKC, Olson M, Pichaud F, Balda MS, Matter K (2017) An apical MRCK-driven morphogenetic pathway controls epithelial polarity. Nat Cell Biol 19(9):1049–1060.

20. Brook JD, McCurrach ME, Harley HG, Buckler AJ, Church D, Aburatani H, Hunter K, Stanton VP, Thirion JP, Hudson T (1992) Molecular basis of myotonic dystrophy: expansion of a trinucleotide (CTG) repeat at the 3’ end of a transcript encoding a protein kinase family member. Cell 68(4):799–808.

21. Mahadevan M, Tsilfidis C, Sabourin L, Shutler G, Amemiya C, Jansen G, Neville C, Narang M, Barceló J, O’Hoy K, LeBlond S, Earle-MacDonald J, de Jong PJ, Wieringa B, Korneluk RG (1992) Myotonic Dystrophy Mutation: an Unstable CTG Repeat in the 30 Untranslated region of the Gene. Science (80-) 255(5049):1253–1255.

22. Fu YH, Pizzuti A, Fenwick RG, King J, Rajnarayan S, Dunne PW, Dubel J, Nasser GA, Ashizawa T, de Jong P, Wieringa B, Korneluk R, Perryman MB, Epstein HF, Caskey CT (1992) An Unstable Triplet Repeat in a Gene Related to Myotonic Muscular Dystrophy. Science (80-) 255(5049):1256–1258.

23. Nishimura T, Takeichi M (2008) Shroom3-mediated recruitment of Rho kinases to the apical cell junctions regulates epithelial and neuroepithelial planar remodeling. Development 135(8):1493–1502.

24. Gaggioli C, Hooper S, Hidalgo-Carcedo C, Grosse R, Marshall JF, Harrington K, Sahai E (2007) Fibroblast-led collective invasion of carcinoma cells with differing roles for RhoGTPases in leading and following cells. Nat Cell Biol 9(12):1392–1400.

25. Unbekandt M, Olson MF (2014) The actin-myosin regulatory MRCK kinases: regulation, biological functions and associations with human cancer. J Mol Med 92(3):217–225.

26. Pearce LR, Komander D, Alessi DR (2010) The nuts and bolts of AGC protein kinases. Nat Rev Mol Cell Biol 11(1):9–22.

27. Yamaguchi H, Kasa M, Amano M, Kaibuchi K, Hakoshima T (2006) Molecular mechanism for the regulation of rho-kinase by dimerization and its inhibition by fasudil. Structure 14(3):589–600.

28. Heikkila T, Wheatley E, Crighton D, Schroder E, Boakes A, Kaye SJ, Mezna M, Pang L, Rushbrooke M, Turnbull A, Olson MF (2011) Co-Crystal Structures of Inhibitors with MRCKβ, a Key Regulator of Tumor Cell Invasion. PLoS One 6(9):e24825.

29. Elkins JM, Amos A, Niesen FH, Pike ACW, Fedorov O, Knapp S (2009) Structure of dystrophia myotonica protein kinase. Protein Sci 18(4):782–91.

30. Matsui T, Amano M, Yamamoto T, Chihara K, Nakafuku M, Ito M, Nakano T, Okawa K, Iwamatsu A, Kaibuchi K (1996) Rho-associated kinase, a novel serine/threonine kinase, as a putative target for small GTP binding protein Rho. EMBO J 15(9):2208–16.

31. Lam LT, Pham YCN, Man N thi, Morris GE (2000) Characterization of a monoclonal antibody panel shows that the myotonic dystrophy protein kinase, DMPK, is expressed almost exclusively in muscle and heart. Hum Mol Genet 9(14):2167–2173.

32. Truebestein L, Elsner DJ, Leonard TA (2016) Made to measure - keeping Rho kinase at a distance. Small GTPases 7(2):82–92.

33. Jumper J, et al. (2021) Highly accurate protein structure prediction with AlphaFold. Nature: 1–7.

34. Senior AW, et al. (2020) Improved protein structure prediction using potentials from deep learning. Nature 577(7792):706–710.

35. Bartual SG, Wei W, Zhou Y, Pravata VM, Fang W, Yan K, Ferenbach AT, Lockhart DEA, van Aalten DMF (2021) The citron homology domain as a scaffold for Rho1 signaling. Proc Natl Acad Sci U S A 118(39). doi:10.1073/pnas.2110298118.

36. Baker NA, Sept D, Joseph S, Holst MJ, McCammon JA (2001) Electrostatics of nanosystems: Application to microtubules and the ribosome. Proc Natl Acad Sci 98(18):10037–10041.

37. Krissinel E, Henrick K (2005) Multiple Alignment of Protein Structures in Three Dimensions. Computational Life Sciences (Springer, Berlin, Heidelberg), pp 67–78.

38. Krissinel E, Henrick K, IUCr (2004) Secondary-structure matching (SSM), a new tool for fast protein structure alignment in three dimensions. Acta Crystallogr Sect D Biol Crystallogr 60(12):2256–2268.

39. Das J, Rahman GM (2014) C1 domains: structure and ligand-binding properties. Chem Rev 114(24):12108–31.

40. Zhang G, Kazanietz MG, Blumberg PM, Hurley JH (1995) Crystal structure of the cys2 activator-binding domain of protein kinase C delta in complex with phorbol ester. Cell 81(6):917–24.

41. Choi SH, Czifra G, Kedei N, Lewin NE, Lazar J, Pu Y, Marquez VE, Blumberg PM (2008) Characterization of the interaction of phorbol esters with the C1 domain of MRCK (myotonic dystrophy kinase-related Cdc42 binding kinase) alpha/beta. J Biol Chem 283(16):10543–9.

42. Okazaki S, Kato R, Uchida Y, Taguchi T, Arai H, Wakatsuki S (2012) Structural basis of the strict phospholipid binding specificity of the pleckstrin homology domain of human evectin-2. Acta Crystallogr D Biol Crystallogr 68(Pt 2):117–23.

43. Saliba A-E, Vonkova I, Ceschia S, Findlay GM, Maeda K, Tischer C, Deghou S, van Noort V, Bork P, Pawson T, Ellenberg J, Gavin A-C (2014) A quantitative liposome microarray to systematically characterize protein-lipid interactions. Nat Methods 11(1):47–50.

44. Saliba A-E, Vonkova I, Deghou S, Ceschia S, Tischer C, Kugler KG, Bork P, Ellenberg J, Gavin A-C (2016) A protocol for the systematic and quantitative measurement of protein–lipid interactions using the liposome-microarray-based assay. Nat Protoc 11(6):1021–1038.

45. Ferguson KM, Lemmon MA, Schlessinger J, Sigler PB (1995) Structure of the high affinity complex of inositol trisphosphate with a phospholipase C pleckstrin homology domain. Cell 83(6):1037–46.

46. Couzens AL, Saridakis V, Scheid MP (2009) The hydrophobic motif of ROCK2 requires association with the N-terminal extension for kinase activity. Biochem J 419(1):141–8.

47. Amano M, Chihara K, Nakamura N, Kaneko T, Matsuura Y, Kaibuchi K (1999) The COOH terminus of Rho-kinase negatively regulates rho-kinase activity. J Biol Chem 274(45):32418–24.

48. Ishizaki T, Naito M, Fujisawa K, Maekawa M, Watanabe N, Saito Y, Narumiya S (1997) p160ROCK, a Rho-associated coiled-coil forming protein kinase, works downstream of Rho and induces focal adhesions. FEBS Lett 404(2-3):118–24.

49. Feng J, Ito M, Kureishi Y, Ichikawa K, Amano M, Isaka N, Okawa K, Iwamatsu A, Kaibuchi K, Hartshorne DJ, Nakano T (1999) Rho-associated Kinase of Chicken Gizzard Smooth Muscle. J Biol Chem 274(6):3744–3752.

50. Tan I, Seow KT, Lim L, Leung T (2001) Intermolecular and Intramolecular Interactions Regulate Catalytic Activity of Myotonic Dystrophy Kinase-Related Cdc42-Binding Kinase⍰α. Mol Cell Biol 21(8):2767–2778.

51. Chugh P, Paluch EK (2018) The actin cortex at a glance. J Cell Sci 131(14). doi:10.1242/jcs.186254.

52. Loo C-S, Chen C-W, Wang P-J, Chen P-Y, Lin S-Y, Khoo K-H, Fenton RA, Knepper MA, Yu M-J (2013) Quantitative apical membrane proteomics reveals vasopressin-induced actin dynamics in collecting duct cells. Proc Natl Acad Sci U S A 110(42):17119–24.

53. Mason FM, Tworoger M, Martin AC (2013) Apical domain polarization localizes act?n-myosin activity to drive ratchet-like apical constriction. Nat Cell Biol 15(8):926–936.

54. Plageman TF, Chauhan BK, Yang C, Jaudon F, Shang X, Zheng Y, Lou M, Debant A, Hildebrand JD, Lang RA (2011) A Trio-RhoA-Shroom3 pathway is required for apical constriction and epithelial invagination. Development 138(23):5177–5188.

55. Mohan S, Rizaldy R, Das D, Bauer RJ, Heroux A, Trakselis MA, Hildebrand JD, VanDemark AP (2012) Structure of Shroom domain 2 reveals a three-segmented coiled-coil required for dimerization, Rock binding, and apical constriction. Mol Biol Cell 23(11):2131–2142.

56. Lee ICJ, Leung T, Tan I (2014) Adaptor protein LRAP25 mediates myotonic dystrophy kinase-related Cdc42-binding kinase (MRCK) regulation of LIMK1 protein in lamellipodial F-actin dynamics. J Biol Chem 289(39):26989–27003.

57. Tan I, Yong J, Dong JM, Lim L, Leung T (2008) A tripartite complex containing MRCK modulates lamellar actomyosin retrograde flow. Cell 135(1):123–36.

58. Ng Y, Tan I, Lim L, Leung T (2004) Expression of the human myotonic dystrophy kinase-related Cdc42-binding kinase gamma is regulated by promoter DNA methylation and Sp1 binding. J Biol Chem 279(33):34156–64.

59. Leung T, Chen XQ, Manser E, Lim L (1996) The p160 RhoA-binding kinase ROK alpha is a member of a kinase family and is involved in the reorganization of the cytoskeleton. Mol Cell Biol 16(10):5313–5327.

60. Garcia P, Ucurum Z, Bucher R, Svergun DI, Huber T, Lustig A, Konarev P V., Marino M, Mayans O (2006) Molecular insights into the self-assembly mechanism of dystrophia myotonica kinase. FASEB J 20(8):1142–51.

61. Lassila JK, Zalatan JG, Herschlag D (2011) Biological phosphoryl-transfer reactions: understanding mechanism and catalysis. Annu Rev Biochem 80(1):669–702.

62. McCoy AJ, Grosse-Kunstleve RW, Adams PD, Winn MD, Storoni LC, Read RJ, IUCr (2007) *Phaser* crystallographic software. J Appl Crystallogr 40(4):658–674.

