## Supplementary Data for "Structure and regulation of the myotonic dystrophy kinase-related Cdc42-binding kinase"

[4]École Polytechnique Fédérale de Lausanne (EPFL), AI 1108, Station 19, CH-1015 Lausanne, Switzerland

[5]University of Geneva, Department of Cell Physiology and Metabolism, CMU Rue Michel-Servet 1 1211 Genève 4, Switzerland

**Supplementary Figures 1-4**

**Supplementary Table 1**

**Figure S1. Full-length MRCK adopts a ROCK-like extended, active conformation.**

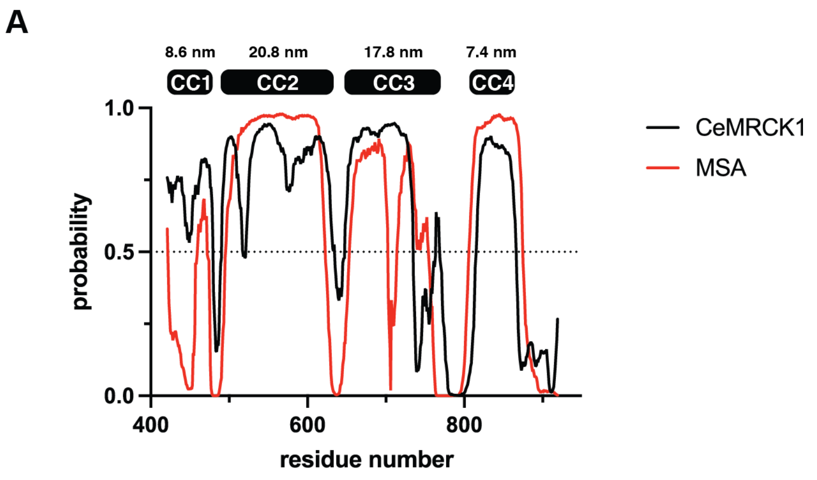


A. DeepCoil2 prediction of the coiled-coil domain of MRCK. Black line: *C. elegans* MRCK1; red line: prediction from multiple sequence alignment (MSA). Estimated lengths of contiguous predicted coiled-coil segments are indicated above the chart.

**Figure S2. Structure and properties of the C1-PH-CNH module of MRCK1.**

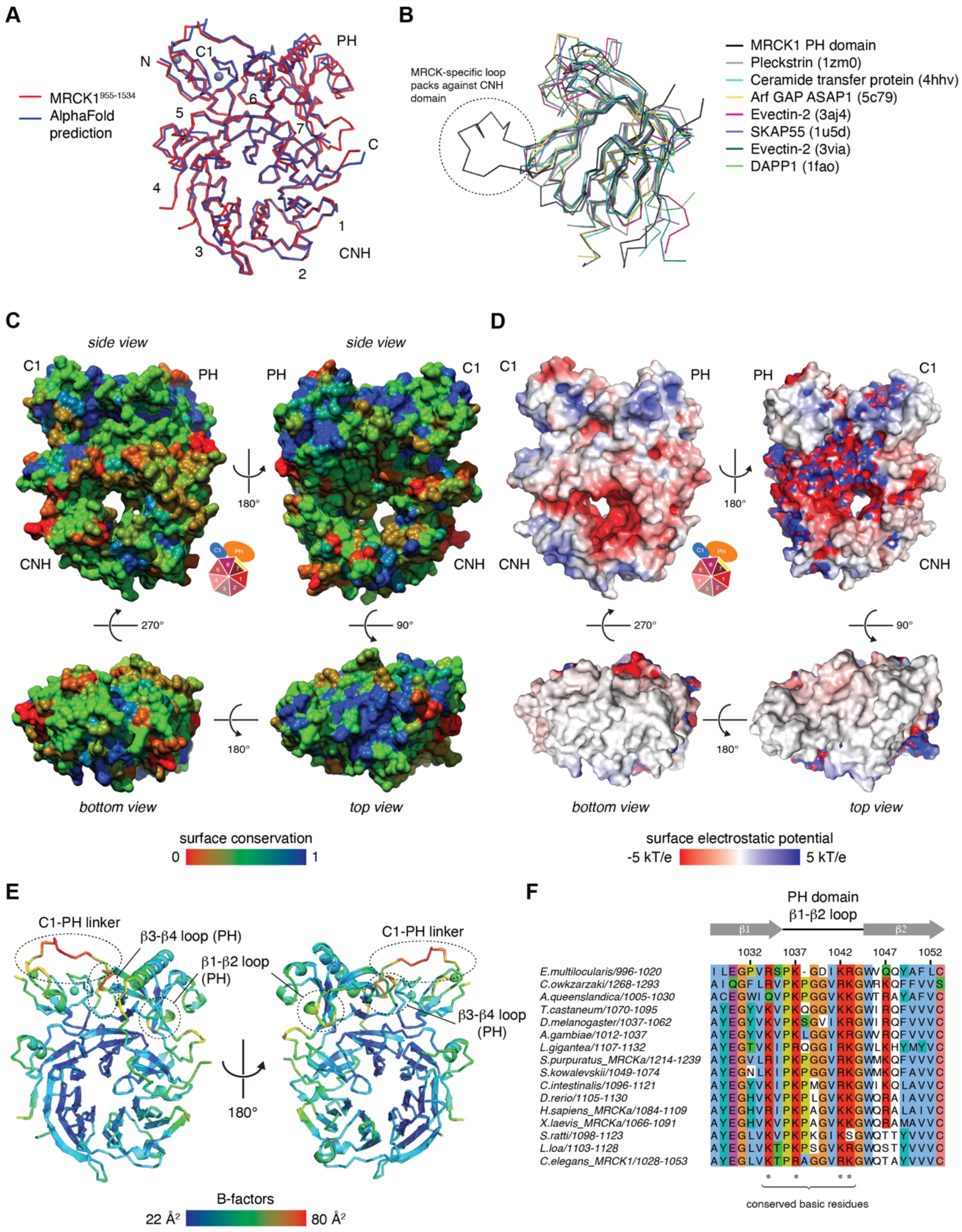


A. Superposition of AlphaFold prediction of MRCK1$^{955\text{-}1534}$ with experimentally determined structure. R.m.s.d. over all atoms = 0.906 Å.

B. Superposition of MRCK1 PH domain with closest structural homologs. Circled:

MRCK-specific loop that packs against the CNH domain.

C. Surface conservation maps of the MRCK1 C1-PH-CNH module. Color gradient: red (least conserved) – green – blue (most conserved).
D. Surface electrostatic potential analysis of MRCK1$^{955\text{-}1534}$.
E. B-factors of final refined model. Regions of higher B-factors include the C1-PH interdomain linker, and the β1-β2 and β3-β4 loops of the PH domain.
F. Sequence alignment of the β1-β2 loop of the PH domain of MRCK. Conserved basic residues in the canonical lipid binding pocket are indicated.

**Figure S3. Membrane-binding properties of the C1-PH-CNH module of MRCK1.**

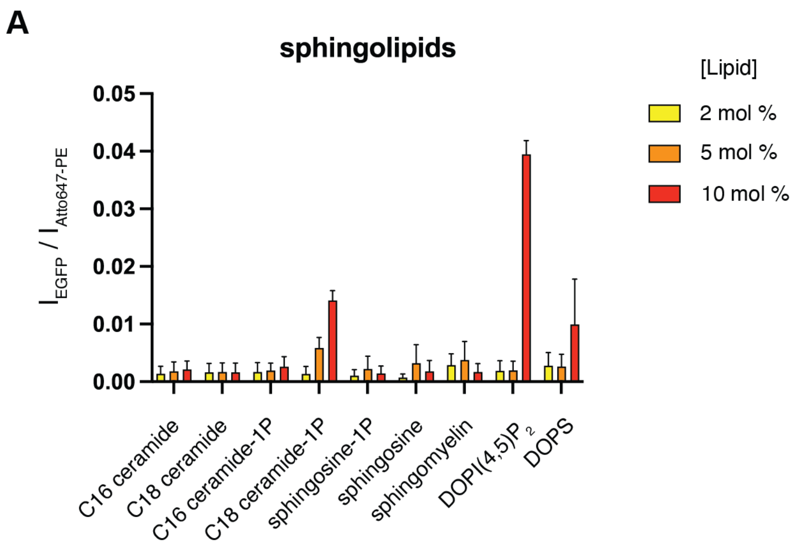


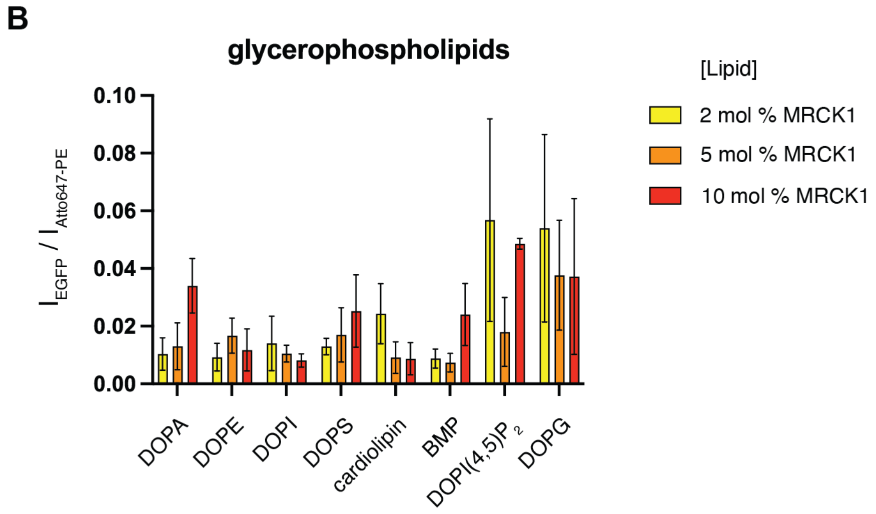


A. Normalized binding of MRCK1$^{955-1534}$-EGFP to sphingolipids. Data derived from LiMA assay. Note that the scale on the y-axis is not the same as in Figure 3B.
B. Normalized binding of MRCK1$^{955-1534}$-EGFP to glycerophospholipids. Data derived from LiMA assay. Note that the scale on the y-axis is not the same as in Figure 3B.

### Figure S4. Model of DMPK kinases as molecular rulers.

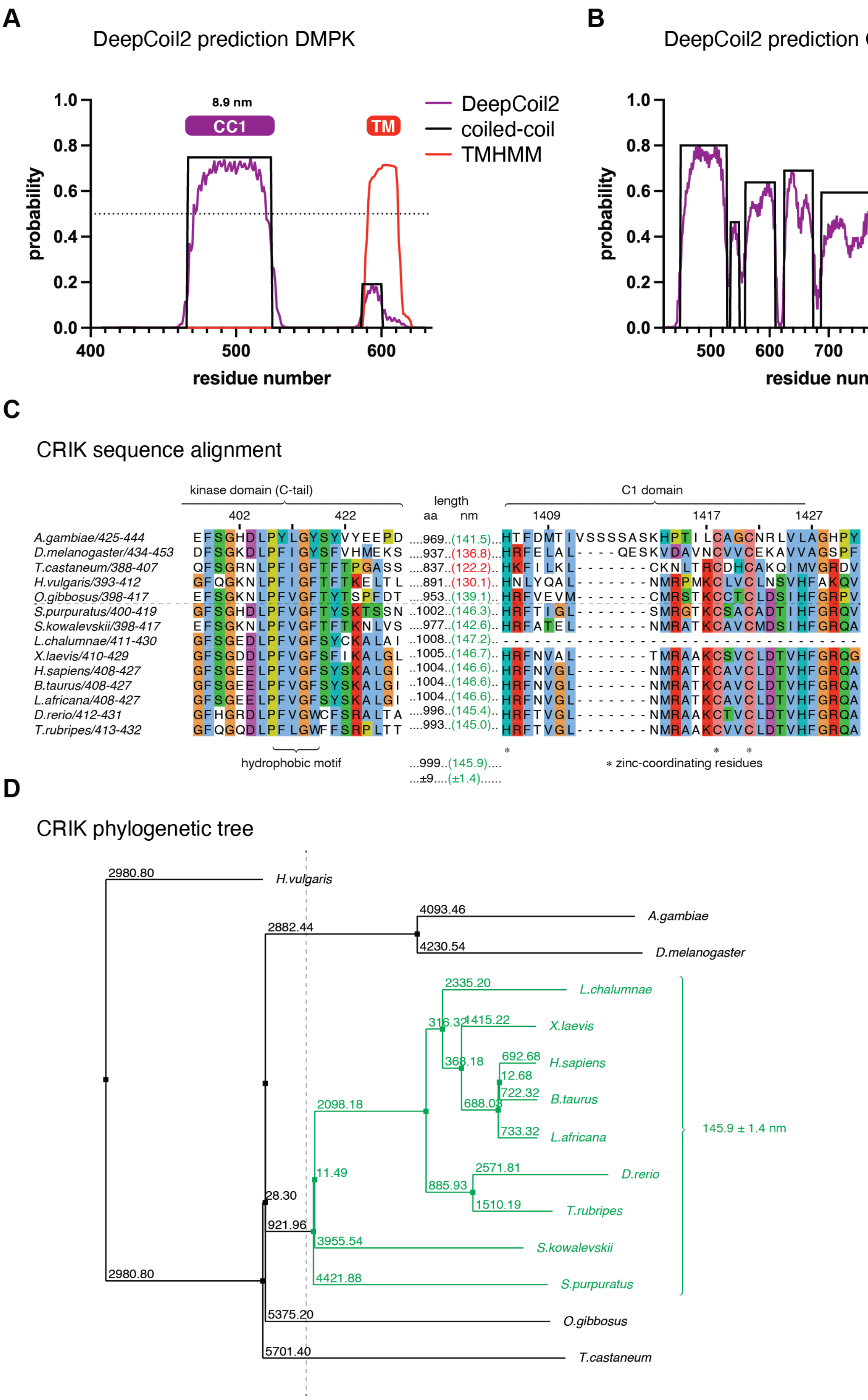

A. DeepCoil2 prediction of the coiled-coil domain (black) and TMHMM prediction of the transmembrane helix (red) of DMPK1.
B. DeepCoil2 prediction of the coiled-coil domain (black) of CRIK.
C. Partial sequence alignment of CRIK orthologs. The length of the intervening sequence between the end of the kinase domain and the beginning of the C1 domain is indicated as number of amino acids (black) and predicted length of a canonical parallel coiled-coil (green). Sequences below the dotted line all belong to the deuterostome branch of the phylogenetic tree which evolved ~550 Myr ago (dashed line).
D. Phylogenetic tree of CRIK orthologs. Deuterostomes are indicated with green branches. The mean predicted length and standard deviation of the coiled-coil domain for deuterostome CRIK orthologs is indicated.

**Supplementary Table 1. Data collection and refinement statistics.**

| | MRCK1 955-1534 |
|---|---|
| Wavelength | 0.9762 |
| Resolution range | 49.07 - 2.14 (2.216 - 2.14) |
| Space group | P 1 |
| Unit cell | 69.142 97.467 131.413 81.3801 77.7664 80.394 |
| Total reflections | 569580 (55034) |
| Unique reflections | 171080 (16283) |
| Multiplicity | 3.3 (3.3) |
| Completeness (%) | 93.96 (89.59) |
| Mean I/sigma(I) | 6.92 (1.05) |
| Wilson B-factor | 35.63 |
| R-merge | 0.1224 (1.167) |
| R-meas | 0.1461 (1.399) |
| R-pim | 0.07886 (0.7611) |
| CC1/2 | 0.993 (0.427) |
| CC* | 0.998 (0.773) |
| Reflections used in refinement | 170148 (16275) |
| Reflections used for R-free | 8480 (803) |
| R-work | 0.1938 (0.3063) |
| R-free | 0.2435 (0.3267) |
| CC(work) | 0.960 (0.720) |
| CC(free) | 0.927 (0.688) |
| Number of non-hydrogen atoms | 21847 |
| macromolecules | 20742 |
| ligands | 10 |
| solvent | 1095 |

| | |
|---|---|
| **Protein residues** | 2628 |
| **RMS(bonds)** | 0.008 |
| **RMS(angles)** | 0.99 |
| **Ramachandran favored (%)** | 96.21 |
| **Ramachandran allowed (%)** | 3.75 |
| **Ramachandran outliers (%)** | 0.04 |
| **Rotamer outliers (%)** | 3.57 |
| **Clashscore** | 6.52 |
| **Average B-factor** | 51.46 |
| **macromolecules** | 51.80 |
| **ligands** | 47.21 |
| **solvent** | 45.09 |
| **Number of TLS groups** | 18 |

Statistics for the highest-resolution shell are shown in parentheses.
